# TGN/EE SNARE protein SYP61 is ubiquitinated and required for carbon/nitrogen-nutrient responses in Arabidopsis

**DOI:** 10.1101/2020.06.03.131094

**Authors:** Yoko Hasegawa, Thais Huarancca Reyes, Tomohiro Uemura, Akari Fujimaki, Yongming Luo, Yoshie Morita, Yasushi Saeki, Anirban Baral, Shugo Maekawa, Shigetaka Yasuda, Koki Mukuta, Yoichiro Fukao, Keiji Tanaka, Akihiko Nakano, Rishikesh P. Bhalerao, Takeo Sato, Junji Yamaguchi

## Abstract

Ubiquitination is a post-translational modification with reversible attachment of the small protein ubiquitin, which is involved in numerous cellular processes including membrane trafficking. For example, ubiquitination of cargo proteins is known to regulate their subcellular dynamics, and plays important roles in plant growth and stress adaptation. However, the regulatory mechanism of the trafficking machinery components remains elusive. Here, we report Arabidopsis *trans-*Golgi network/early endosome (TGN/EE) localized soluble *N*-ethylmaleimide sensitive factor attachment protein receptor (SNARE) protein SYP61 as a novel ubiquitination target of a membrane localized ubiquitin ligase ATL31. SYP61 is a key component of membrane trafficking in Arabidopsis. SYP61 was ubiquitinated with K63-linked chain by ATL31 *in vitro* and in plants. The knockdown mutants of *SYP61* were hypersensitive to the disrupted carbon (C)/nitrogen (N)-nutrient stress, suggesting its critical role in plant homeostasis in response to nutrients. We also found the ubiquitination status of SYP61 is affected by C/N-nutrient availability. These results provided possibility that ubiquitination of SNARE protein has important role in plant physiology.

## Introduction

Ubiquitination plays critical roles in regulating various proteins and enables flexible responses toward the changing environment(Hershko & Ciechanover, 1998; Oh, Akopian, & Rape, 2018). Ubiquitin is an 8.5 kDa protein, which is attached to lysine (K) residues of target proteins by ubiquitin ligases. As ubiquitin can form chain through its own seven K and one N-terminal methionine residues, there are several types of topologically different ubiquitin chains, which modulate distinct cellular processes(Oh et al., 2018). For example, proteins modified by K48 and K11-linked ubiquitin chains undergo proteasomal degradation, whereas K63-linked ubiquitinated proteins are involved in several processes, including vacuolar targeting, endocytosis, DNA repair and signal activation(Callis, 2014; Isono & Kalinowska, 2017; Oh et al., 2018; Romero-Barrios & Vert, 2018). Protein ubiquitination is involved in regulating the membrane trafficking of cargo proteins during plant responses to environmental stresses. For example, multiple mono-ubiquitination of the plant metal transporter IRT1 is necessary for its constitutive turnover from the plasma membrane to the TGN/EE, with extension of ubiquitination into K63-linked chains in the presence of excess amounts of non-iron metal ions leading to vacuolar degradation(Barberon et al., 2011; Dubeaux, Neveu, Zelazny, & Vert, 2018). Ubiquitination of other plasma membrane localized proteins, including brassinosteroid receptor BRI1, and auxin efflux carrier PIN2, also triggers endocytosis and vacuolar targeting(Korbei et al., 2013; Leitner et al., 2012; Martins et al., 2015; Zhou et al., 2018). Ubiquitination of boron transporter BOR1 was reported to be necessary for its vacuolar degradation under high concentration of boron(Kasai, Takano, Miwa, Toyoda, & Fujiwara, 2011), and mono-ubiquitination of a membrane associated receptor-like cytoplasmic kinase BIK1 was recently reported to trigger its endocytosis and plays important role for activation of plant immunity(Ma et al., 2020). Deubiquitination reactions are also important for the cargo protein trafficking via the ESCRT system(Isono & Kalinowska, 2017). Although these previous studies showed trafficking regulation of cargo proteins, less is known about the ubiquitination of the trafficking machinery itself.

SNARE proteins are key regulators of membrane trafficking, which are conserved among eukaryotes(Jahn & Scheller, 2006; Lipka, Kwon, & Panstruga, 2007; Wickner & Schekman, 2008). They mediate vesicle fusion by forming SNARE complexes. The Arabidopsis genome encodes more than 60 SNARE proteins(Lipka et al., 2007; A. Sanderfoot, 2007; Anton A Sanderfoot, Assaad, & Raikhel, 2000; Uemura et al., 2004). Each SNARE protein localizes to specific membranes, and has specific SNARE partners in the cells (Fujiwara et al., 2014; Jahn & Scheller, 2006; Pratelli, Sutter, & Blatt, 2004; Uemura et al., 2004). Some SNARE proteins are reported to participate in several SNARE complexes and mediate different trafficking pathways. For example, a TGN/EE localized SNARE protein, SYNTAXIN OF PLANTS 61 (SYP61) is reported to mediate vacuolar trafficking with SYP41, VPS TEN INTERACTING 12 (VTI12), and a Sec1/Munc18 (SM) family protein, VACUOLAR PROTEIN SORTING 45 (VPS45)(Kim & Bassham, 2011; Zouhar, Rojo, & Bassham, 2009), and retrograde and anterograde trafficking of aquaporin PLANT PLASMA MEMBRANE INTRINSIC PROTEINS 2;7 (PIP2;7), possibly with SYP121(Hachez et al., 2014). SYP61 is also reported to mediate exocytotic trafficking of cell wall components(Drakakaki et al., 2012; Gendre et al., 2013). SNARE proteins play important roles in plant physiological responses. For *SYP61*, the knockdown mutant was previously reported to be hypersensitive to the salt and osmotic stresses(Zhu et al., 2002).

Sugar (carbon, C) and nitrogen (N) are essential components of organisms, and their relative availability, C/N-nutrient balance, affects many aspects of plant physiology. Disrupted high C/low N nutrient conditions inhibit early post-germination growth and promote the progression of senescence in Arabidopsis plants(Aoyama et al., 2014; Coruzzi & Zhou, 2001; Martin, Oswald, & Graham, 2002). We previously reported that a membrane localized ubiquitin ligase ATL31 regulates C/N-nutrient response in Arabidopsis(Sato et al., 2009; Yasuda, Aoyama, Hasegawa, Sato, & Yamaguchi, 2017). The knockout mutant of *ATL31* is hypersensitive, and the overexpressor is insensitive to the high C/low N-nutrient stress. This function of ATL31 is depending on both of the membrane localization and the ubiquitin ligase activity(Sato et al., 2009).

In this study, we identified SYP61 as a direct target of ATL31. We also demonstrate that SYP61 is involved in plant C/N-nutrient stress responses, and the ubiquitination status was affected by C/N-nutrient availability.

## Results

### ATL31 localizes to the plasma membrane and endosomal compartments

We previously showed that the N-terminal transmembrane domain of ATL31 is required for its function in the C/N-nutrient stress responses(Sato et al., 2009). Therefore, we performed subcellular localization analysis of ATL31. In the root tip cells of Arabidopsis plants expressing green fluorescent protein-tagged ATL31 (ATL31-GFP), ATL31-GFP signals were detected at the intracellular punctate structures and the plasma-membrane under confocal laser microscopy (Fig. 1A). About 56% of the root epidermal cells showed ATL31-GFP in intracellular dot-like structures alone, and 44% showed signals in the plasma membranes and dot-like structures (Fig. 1B). The identity of these dot-like structures was further determined by co-localization analysis with monomeric red fluorescent protein (mRFP)-tagged endomembrane organelle markers. Transient expression in Arabidopsis protoplast cells showed that ATL31-GFP signals were detected in mRFP-SYP61 (TGN/EE marker) and mRFP-VAMP727 (late endosome (LE) marker) positive structures, but not in mRFP-tagged ST (ST-mRFP, Golgi marker) positive structures (Supplementary Fig. S1). The subcellular localization was also confirmed in transgenic Arabidopsis plants co-expressing ATL31-GFP and each RFP-tagged marker. ATL31-GFP co-localized with the TGN/EE markers mRFP-SYP43, mRFP-SYP61 and VHAa1-mRFP; and the LE markers mRFP-ARA7 and tagRFP-VAMP727 (Fig. 1C). These results demonstrate that ATL31 localizes to the plasma membrane and multiple endosomal compartments positive for TGN/EE and LE markers.

**Fig. 1.**
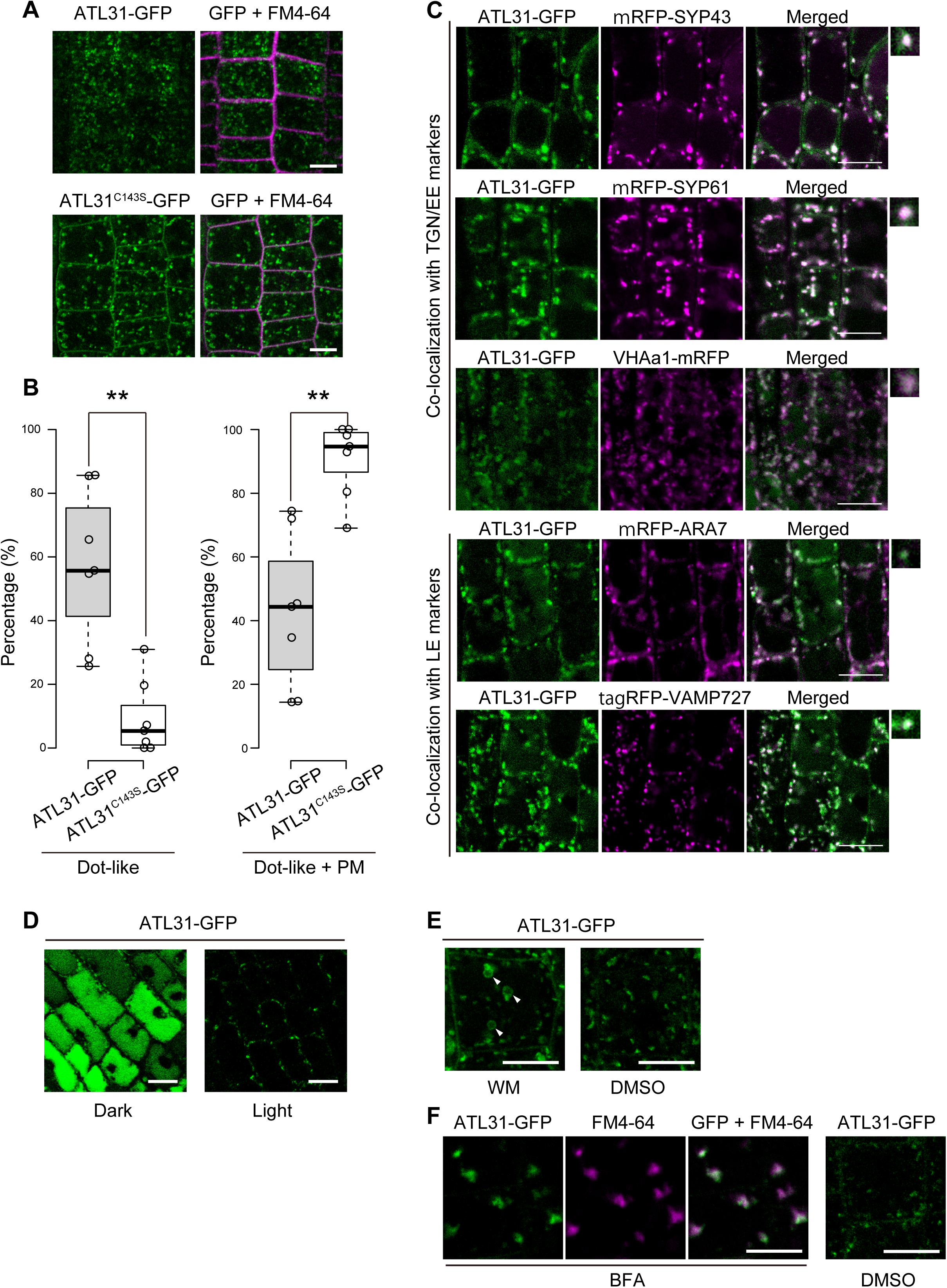
Subcellular localization analysis of ATL31. **(A)** Representative confocal images of Arabidopsis roots expressing ATL31-GFP or ATL31^C143S^-GFP. Plasma membrane stained with FM4-64 is shown in magenta. Bars = 10 μm. **(B)** Percentage of cells in a field shows fluorescence in dot like structures alone (left), or dot like structures and plasma membranes (PM) (right). n > 40 cells in a root for each experiment, 7 biological repeats. Statistical significance was determined by two-tailed Welch’ s *t-*test (\*\**P*<0.01). **(C)** Representative confocal images of Arabidopsis roots co-expressing ATL31-GFP with mRFP-SYP43 (TGN/EE), mRFP-SYP61 (TGN/EE), VHAa1-mRFP (TGN/EE), mRFP-ARA7 (LE), and tagRFP-VAMP727 (LE). Right panels show enlarged view of the dot-like structures. LE, late endosome. Bars = 10 μm. **(D)** Representative confocal images of Arabidopsis roots expressing ATL31-GFP after light or dark treatment for 10 hours. Plasma membranes stained with FM4-64 are shown in magenta. Bars = 10 μm. **(E)** Representative confocal images of Arabidopsis roots expressing ATL31-GFP treated with 33 μM Wortmannin (WM) or DMSO for 1 h. Arrowheads indicate the ring like structures. Bars = 10 μm. **(F)** Representative confocal images of Arabidopsis roots expressing ATL31-GFP treated with 50 μM Brefeldin A (BFA) or DMSO for 30 min. Cells were stained with FM4-64 15 min before BFA treatment. Bars = 10 μm.

Membrane localized proteins often undergo vacuolar degradation, which can be visualized by the signal of the tagged GFP in the vacuole after dark treatment(Tamura et al., 2003). After 10 h dark treatment, GFP fluorescence was observed in vacuoles and in intracellular punctuate structures (Fig. 1D), suggesting that ATL31-GFP is subjected to vacuolar degradation in Arabidopsis cells. We also tested the sensitivity to commonly used membrane traffic inhibitors wortmannin (WM) and Brefeldin A (BFA). WM in an inhibitor of phosphoinositide 3-kinase (PI3K) and PI4K, which causes swelling and vacuolization of late endosomal compartments(Jaillais, Fobis-Loisy, Miège, & Gaude, 2008; Wang, Cai, Miao, Lam, & Jiang, 2009). BFA inhibits the BFA-sensitive ADP-ribosylation factors guanine nucleotide exchange factors (Arf-GEFs) and generates BFA bodies, large aggregations of endosomal compartments(Dettmer, Hong-hermesdorf, Stierhof, & Schumacher, 2006; Geldner et al., 2003; Grebe et al., 2003; Robinson, Jiang, & Schumacher, 2008). Upon WM treatment, ATL31-GFP was observed as small ring-like structures, a finding which was not observed after mock treatment (Fig. 1E). BFA induced the aggregation of ATL31-GFP with fluorescence overlapping FM4-64 containing BFA bodies (Fig. 1F). Altogether, we confirmed that ATL31 localizes to the TGN/EE, LE and PM, and degraded in the vacuole.

### Ubiquitination activity of ATL31 affects its subcellular localization pattern

Because the function of ATL31 in C/N stress responses is dependent on its ubiquitination activity on the RING domain(Sato et al., 2009), we also examined the subcellular localization of the inactive form of ATL31 with a point-mutation at the conserved cysteine-143 residue in the RING domain to serine (ATL31^C143S^)(Sato et al., 2009). Confocal laser microscopy of cells bearing GFP-tagged ATL31^C143S^ (ATL31^C143S^-GFP) showed a localization pattern different from that of intact ATL31-GFP. About 90% of cells showed ATL31^C143S^-GFP signals at both the plasma-membrane and the intracellular dot-like structures, with only 10% showing ATL31^C143S^-GFP signals in the dot-like structures alone (Fig. 1B), suggesting that ATL31 localization is modulated by its ubiquitination activity.

### ATL31 can catalyze several different types of ubiquitination

To further understand the ubiquitin signal mediated by ATL31, we determined the chain type of ubiquitination catalyzed by ATL31. *In vitro* ubiquitination reaction was performed with recombinant ATL31 (MBP-ATL31), and the chain type was quantitatively determined by parallel reaction monitoring (PRM), a targeted tandem mass spectrometry, with Ub-absolute quantification (Ub-AQUA) peptides(Kirkpatrick, Gerber, & Gygi, 2005; Tsuchiya et al., 2017). Mass spectrometry showed that ATL31 catalyzed multiple types of ubiquitin chains, mainly those with K11, K48 and K63 linkages (Fig. 2A). Immunoprecipitation and immunoblotting analyses of ATL31-FLAG expressed in *N. benthamiana* leaves also demonstrated that ATL31 catalyzed K48 and K63 linked ubiquitin chains in plant cells (Fig. 2B).

**Fig. 2.**
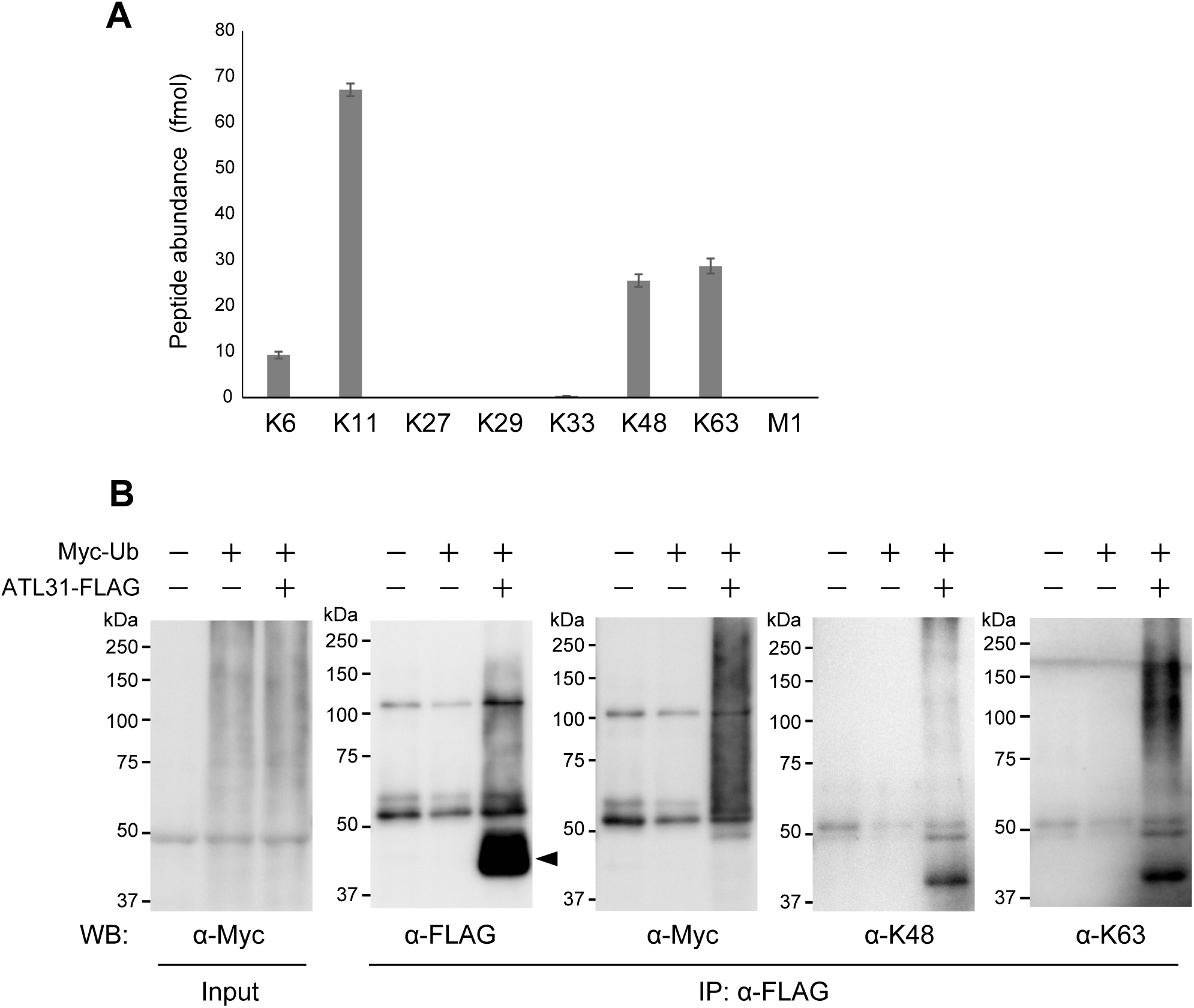
ATL31 catalyzes several types of ubiquitination. **(A)** Quantitative analysis of ubiquitin linkages catalyzed by ATL31 *in vitro*. Purified MBP-ATL31 protein was incubated with E1, E2, ubiquitin and ATP for 10 min. The amount of ubiquitin chain was quantified by the Ub-AQUA/PRM system. Data are means ± s.d. (n = 3). **(B)** ATL31 can form K48- and K63-linked ubiquitin chains in plants. ATL31-FLAG (43.2 kDa, indicated by arrowhead) and Myc-Ub proteins were transiently expressed in *N. benthamiana* leaves. Proteins were immunoprecipitated with anti-FLAG beads, and detected by anti-FLAG, anti-Myc, anti-K48- (Apu2) or anti-K63- (Apu3) linked ubiquitin specific antibodies.

### ATL31 physically interacts with TGN/EE SNARE SYP61

In a proteomic analysis of Arabidopsis TGN/EE SNARE protein SYP61 compartment, ATL6, the highest homologue of ATL31 was identified(Drakakaki et al., 2012). SYP61 is a Qc-SNARE with N-terminal three helical domains and a C-terminal transmembrane region(A. A. Sanderfoot, Kovaleva, Bassham, & Raikhel, 2001; Uemura et al., 2004), and is known to mediate the secretion of cell wall components(Drakakaki et al., 2012; Wilkop et al., 2019). Our previous report suggested that ATL31 is also involved in the accumulation of cell wall components(Maekawa et al., 2014). Considering the functional relevance, we attempted to evaluate the interaction between ATL31 and SYP61. FLAG-tagged intact ATL31, ATL31^C143S^ and GFP-tagged SYP61 were transiently co-expressed in *N. benthamiana* leaves and subjected to co-immunoprecipitation assay. SYP61 was co-immunoprecipitated with both intact ATL31 and ATL31^C143S^ (Fig. 3A). Physical interactions were confirmed by split ubiquitin yeast two-hybrid analysis (Fig. 3B). In addition, immunoblotting with anti-SYP61 antibody detected endogenous SYP61 in the immunoprecipitate of ATL31-GFP from transgenic Arabidopsis plants (Fig. 3C). These results demonstrated that ATL31 physically interacts with SYP61 in plants.

**Fig. 3.**
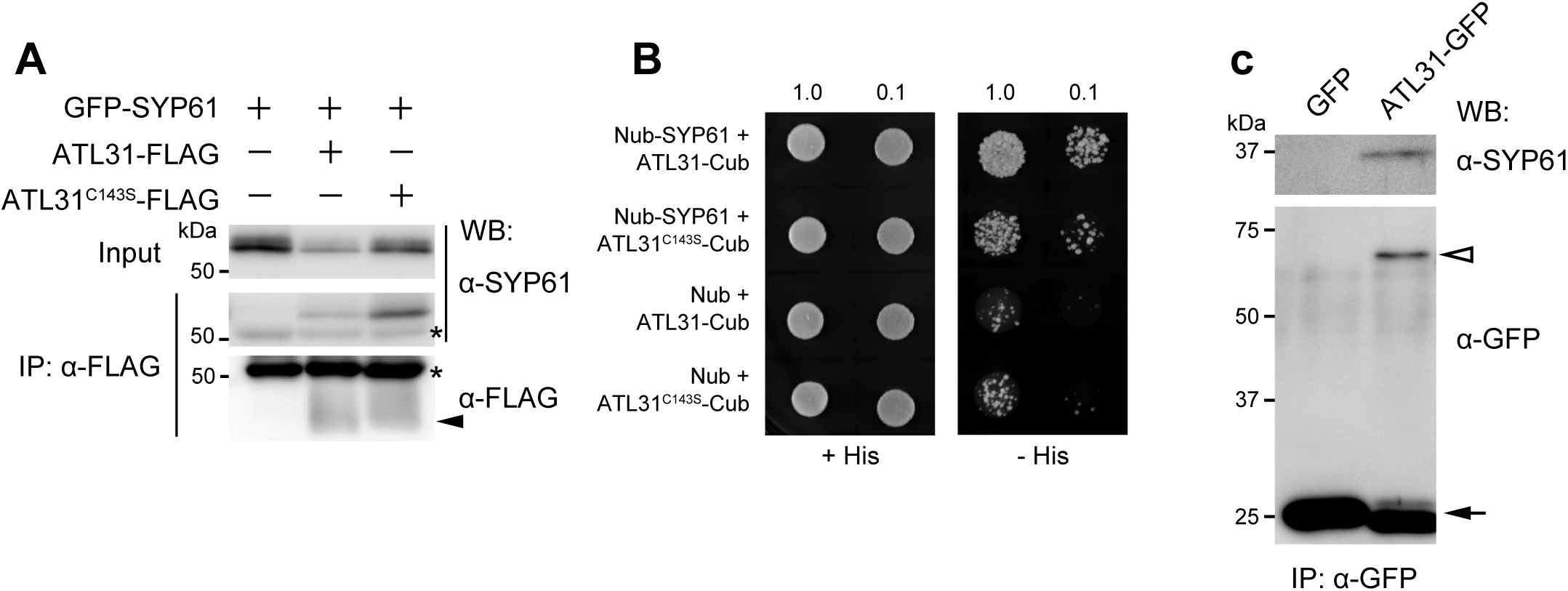
ATL31 interacts with SYP61. **(A)** Co-immunoprecipitation of GFP-SYP61 (56.2 kDa) with ATL31-FLAG or ATL31^C143S^-FLAG (43.2 kDa, indicated by arrowhead). Proteins were expressed in *N. benthamiana* leaves, and immunoprecipitated with anti-FLAG antibody beads, followed by immunoblotting with anti-SYP61 and anti-FLAG antibodies. Asterisks, nonspesific bands and IgG heavy chain. **(B)** Split ubiquitin yeast-two hybrid assays of ATL31-Cub and ATL31^C143S^-Cub with Nub-SYP61 or empty vector. Yeast cultures were diluted to 1 or 10 times (indicated 1.0 or 0.1 each), and grown on solid medium with or without histidine (His). Cub, C-terminal half of ubiquitin; Nub, N-terminal half of ubiquitin. **(C)** Co-immunoprecipitation of endogenous SYP61 with ATL31-GFP in Arabidopsis. Proteins were extracted from Arabidopsis plants expressing GFP (27 kDa, indicated by arrow) or ATL31-GFP (71 kDa, indicated by open arrowhead), and subjected to immunoprecipitation with anti-GFP antibody beads, followed by immunoblotting with anti-SYP61 and anti-GFP antibodies.

### SYP61 protein is ubiquitinated in plant cells

Because ATL31 is a ubiquitin ligase, we asked the possibility of ubiquitin modification on SYP61. First, we extracted proteins from transgenic Arabidopsis plants expressing GFP-fused SYP61 under the control of native promoter (*pSYP61:GFP-SYP61*). GFP-SYP61 was purified by immunoprecipitation in the absence or presence of inhibitors of deubiquitination enzymes (DUBi). Immunoblotting with anti-GFP antibody detected a signal corresponding to intact GFP-SYP61, whereas immunoblotting with anti-ubiquitin antibody yielded three sharp bands of higher molecular weight, which were enhanced by DUBi treatment (Fig. 4A). We also detected ubiquitination of FLAG-tag fused SYP61 (FLAG-SYP61), but not of FLAG-GFP protein in *N. benthamiana* leaves (Supplementary Fig. S2). Based on their molecular weights, the bands detected with ubiquitin antibody were likely ubiquitinated SYP61, not GFP- or FLAG-tagged proteins or the other interactors. To further assess the biochemical significance of SYP61 ubiquitination, we analyzed the ubiquitin chain type. The second band of ubiquitinated SYP61 was also bound by antibody specific for K63-linked, but not K48 linked, ubiquitination (Fig. 4B). These findings revealed that SYP61 was ubiquitinated in plants, and that some pools of SYP61 are modified by K63-type ubiquitination.

**Fig. 4.**
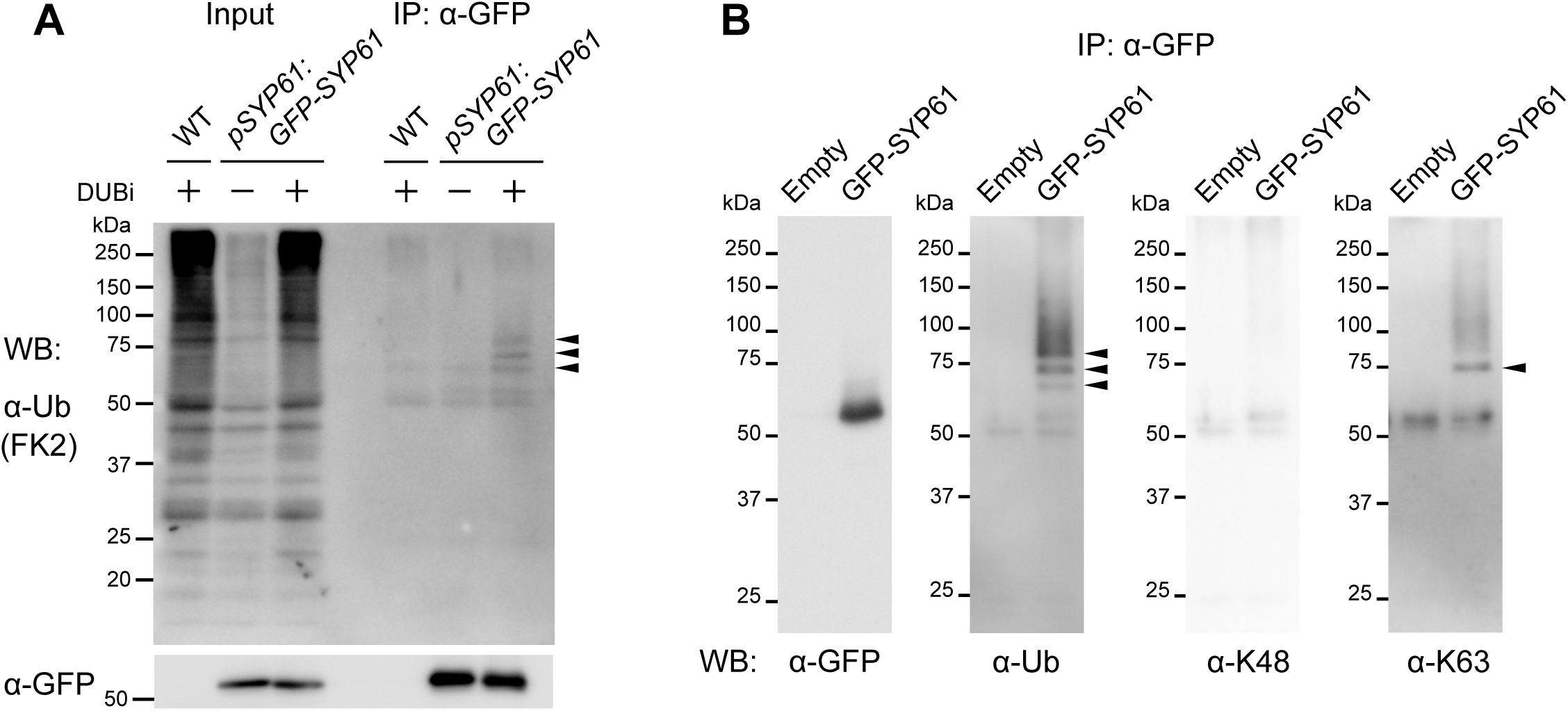
SYP61 is ubiquitinated in plants. **(A)** Proteins were extracted from wild type (WT) or GFP-SYP61 (54.7 kDa) expressing Arabidopsis plants (*pSYP61:GFP-SYP61*) in the presence or absence of deubiquitination enzyme inhibitors (DUBi). Proteins were immunoprecipitated with anti-GFP antibody beads, and detected by anti-ubiquitin (FK2) or anti-GFP antibodies. Arrowheads indicate ubiquitinated GFP-SYP61. **(B)** GFP-SYP61 (56.2 kDa) was transiently expressed in *N. benthamiana* leaves. Extracted proteins were immunoprecipitated with anti-GFP antibody beads, and detected with anti-GFP, anti-ubiquitin (FK2), anti-K48-linked ubiquitin (Apu2), and anti-K63-linked ubiquitin (Apu3) antibodies. Arrowheads indicate ubiquitinated GFP-SYP61.

### SYP61 protein is ubiquitinated by ATL31 *in vitro* and in plant cells

To examine the direct involvement of ATL31 in SYP61 ubiquitination, we performed *in vitro* ubiquitination assays with MBP-tag fused ATL31 (MBP-ATL31) and GST-tag fused SYP61 (GST-SYP61) recombinant proteins. Incubation of GST-SYP61 with MBP-ATL31 for 1 or 3 hours, followed by immunoblotting with either anti-GST or anti-ubiquitin antibody, showed time-associated shifting of the GST-SYP61 bands (Fig. 5A). This ubiquitination signal was not detected after incubation with GST protein, indicating that SYP61 was ubiquitinated by ATL31 *in vitro*. The molecular size of the bands suggested that SYP61 was ubiquitinated with at least one (mono-ubiquitin) and two (multiple mono- or di-ubiquitin) molecules. The two-ubiquitin-bound SYP61 was also detected with anti-K63 ubiquitination antibody (Fig. 5A), indicating that ATL31 catalyzes the attachment of a K63-linked ubiquitin chain to SYP61. Mass spectrometry showed that five lysine residues on SYP61, four on the SNARE domain, were ubiquitinated by ATL31 (Fig. 5B, Supplementary Table S1). We also confirmed that ATL31 ubiquitinates SYP61 in plant cells. GFP-SYP61 was expressed alone or with 3xFLAG tagged ATL31 (ATL31-3xFLAG) or ATL31^C143S^ (ATL31^C143S^-3xFLAG) in *N. benthamiana* leaves by agroinfiltration. As a result, co-expression of ATL31-3xFLAG enhanced signal of the ubiquitinated and the K63-chain ubiquitinated SYP61 (Fig. 5C). This signal enhancement was not observed with the inactive form of ATL31, indicating ATL31 enhanced SYP61 ubiquitination through its ubiquitination activity.

**Fig. 5.**
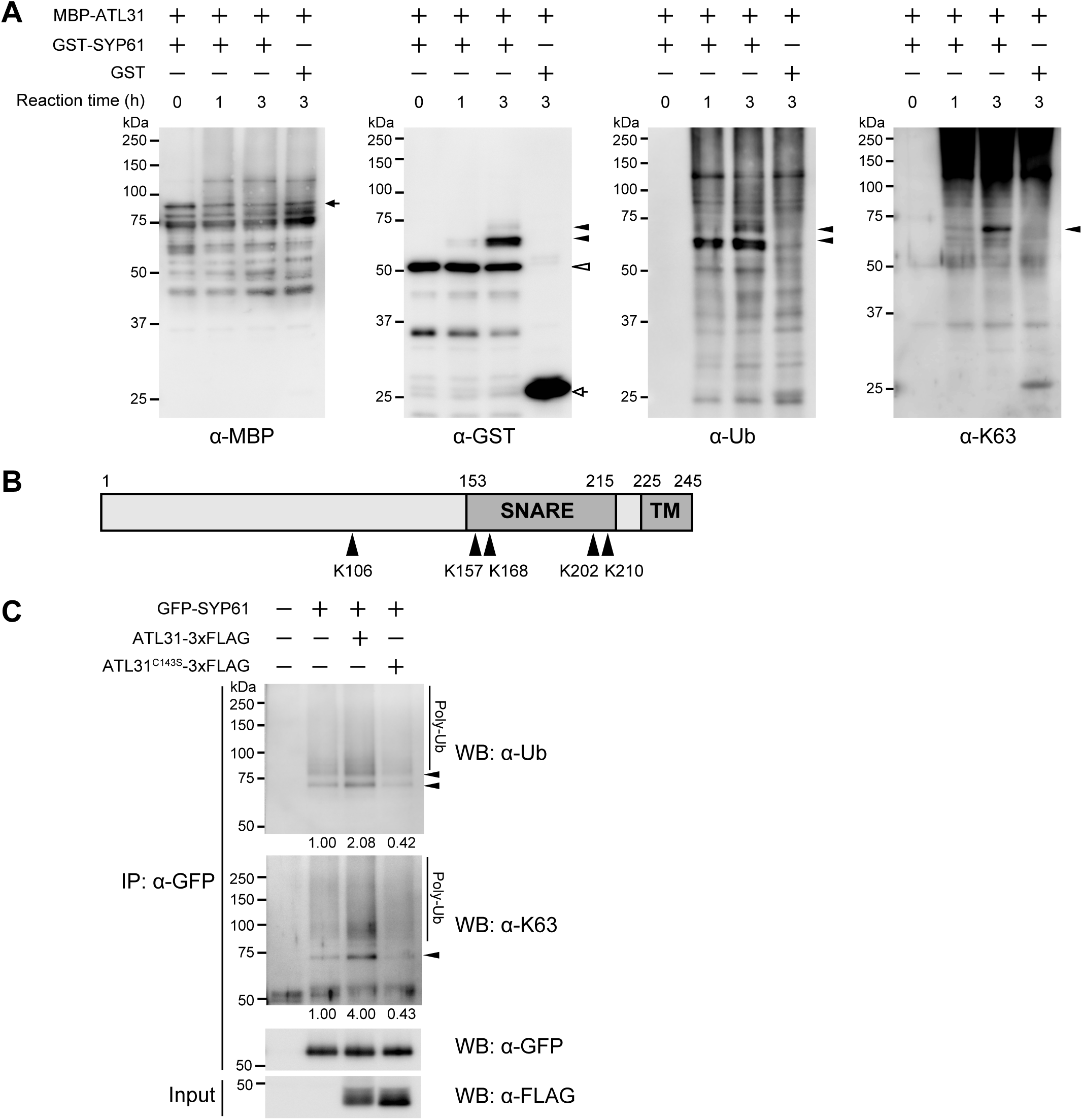
ATL31 ubiquitinates SYP61 *in vitro* and *in planta.* **(A, B)** *In vitro* ubiquitination assay of SYP61 by ATL31. GST-SYP61 or GST was incubated with MBP-ATL31 in the presence of ubiquitin, E1, E2 and ATP for the indicated time. **(A)** Immunoblotting analysis with anti-MBP, anti-GST, anti-ubiquitin (FK2) and anti-K63-linked ubiquitin (Apu3) antibodies. Closed arrow, MBP-ATL31 (80.9 kDa); open arrow, GST (27.9 kDa); open arrowhead, GST-SYP61 (52.6 kDa); closed arrowheads, ubiquitinated SYP61. **(B)** Schematic diagram of the primary structure of SYP61 protein. Arrowheads indicate the ATL31-catalyzed ubiquitination sites of SYP61, as determined by mass spectrometry analysis. Numbers are showing the amino acid positions. SNARE, SNARE domain; TM, transmembrane domain. **(C)** GFP-SYP61 (56.2 kDa) was transiently expressed alone or with ATL31-3xFLAG or ATL31^C143S^-3xFLAG (44.9 kDa) in *N. benthamiana* leaves. Extracted proteins were immunoprecipitated with anti-GFP antibody beads, and detected with anti-GFP, anti-ubiquitin (FK2), anti-K48-linked ubiquitin (Apu2), and anti-K63-linked ubiquitin (Apu3) antibodies. Arrowheads indicate ubiquitinated GFP-SYP61. The numbers are showing the relative intensity of the di-Ub bands (the second band from the top in anti-ubiquitin) normalized by the band intensity of GFP-SYP61.

### TGN/EE SNARE SYP61 is required for plant response to disrupted C/N-nutrient balance

As ATL31 plays an essential role in the response to disrupted carbon/nitrogen (C/N)-nutrient availability, we examined whether TGN/EE SNARE SYP61 is also involved in this response. Post-germination growth of Arabidopsis is inhibited by an increase in sugar with limited nitrogen availability (high C/low N), and these plants exhibit purple pigmentation due to the accumulation of anthocyanin(Martin et al., 2002),(Sato et al., 2009). We examined the sensitivity of *SYP61* knockdown mutants (*syp61 amiRNA*) to high C/low N-nutrient stress conditions. The knockdown mutant plants were generated using an artificial *SYP61* microRNA, and we confirmed that the expression level of *SYP61* was markedly decreased (Supplementary Fig. S3). Two independent lines of the *syp61 amiRNA* knockdown mutants showed significant hypersensitive phenotypes to 150 mM glucose (Glc)/0.3 mM N conditions with no green cotyledons, whereas 65% of wild type seedlings showed expanded green cotyledons (Fig. 6A-6B). We also tested *osm1*, a T-DNA inserted *syp61* mutant previously reported(Zhu et al., 2002). Because this mutant is established on a C24 accession background, which was more resistant to high C/low N-nutrient stress, we examined the *osm1* phenotype in medium containing more glucose than the medium used for Col-0 background mutants. In line with the result in Col-0 plants, the percentage of green cotyledons was significantly lower for *osm1* (10%) than for C24 wild-type (27%) under 300 mM Glc/0.3 mM N stress conditions (Fig. 6C-6D), indicating that the *osm1* mutant was also hypersensitive to high C/low N-nutrient conditions. As all wild-type and the *syp61* mutant seedlings showed green cotyledons in mannitol medium, it was confirmed that the effects of C/N-nutrient stress are distinct from that of osmotic stress (Fig. 6C-6D). These results demonstrated that SYP61 plays an essential role in plant adaptation to C/N-nutrient conditions.

**Fig. 6.**
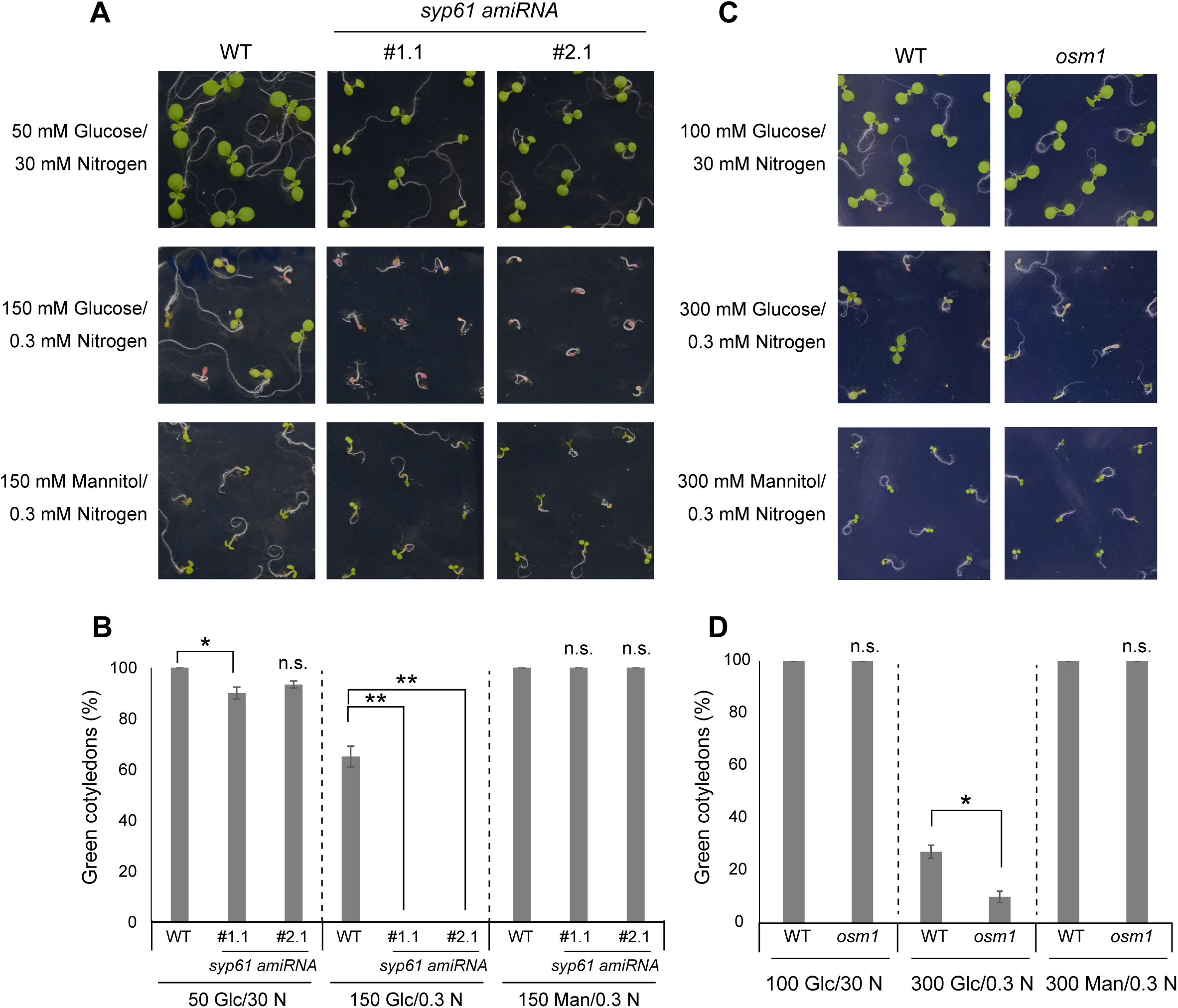
*syp61* mutants are hypersensitive to C/N-nutrient stress. C/N-nutrient responses during post-germination growth. Seedlings were grown on medium containing the indicated concentrations of glucose, nitrogen and mannitol (osmotic control). **(A)** Representative images of WT (Col-0) and *syp61* knockdown mutant (*syp61 amiRNA*) seedlings grown for 8 days on 50 mM glucose/30 mM nitrogen, 150 mM glucose/0.3 mM nitrogen or 150 mM mannitol/0.3 mM nitrogen. **(B)** Percentage of seedlings with green cotyledons after growth for 8 days. Glc, glucose; N, nitrogen; Man, mannitol; unit, mM. WT, wild type. Data are means ± s.e.m. (n = 3). Statistical significance was determined by two-tailed Dunnett’s test. (\**P*<0.05, \*\**P*<0.01). **(C)** Representative images of WT (C24) and *osm1* seedlings grown for 10 days on 100 mM glucose/30 mM nitrogen, or 14 days on 300 mM glucose/0.3 mM nitrogen or 300 mM mannitol/0.3 mM nitrogen. **(D)** Percentage of seedlings with green cotyledons after growth for 14 days. Glc, glucose; N, nitrogen; Man, mannitol; unit, mM. WT, wild type. Data are means ± s.e.m. (n = 3). Statistical significance was determined by two-tailed Welch’s *t-*test. (\**P*<0.05).

### Ubiquitination of SYP61 is affected by C/N-nutrient availability

To assess the relevance of SYP61 ubiquitination in the plant C/N-nutrient responses, we analyzed the ubiquitination status of SYP61 in various C/N-nutrient conditions (Fig. 7). The Arabidopsis transgenic plant *pSYP61:GFP-SYP61* was grown in liquid medium containing 100 mM Glc/30 mM N for 10 days, and treated with 6 different media containing 0, 100 or 200 mM Glc and 30 or 0.3 mM N for 3 hours. Interestingly, the signal of the ubiquitinated SYP61 was the highest in the 0 mM Glc/30 mM N treated sample. The difference of the signal of K63-linked chain was even more significant. The signal was higher in the 0 mM Glc treated samples than the other glucose conditions with the same nitrogen status. Comparing the two 0 mM Glc treated samples with different N conditions, the one with 30 mM N showed higher band intensity than the one with 0.3 mM N. Because the ubiquitination signals were not decreased by the addition of the mannitol instead of glucose, it was confirmed that the observed difference was not due to the osmotic conditions (Supplementary Fig. S4). Taken together, both of the glucose and nitrogen conditions affected the ubiquitination status of SYP61.

**Fig. 7.**
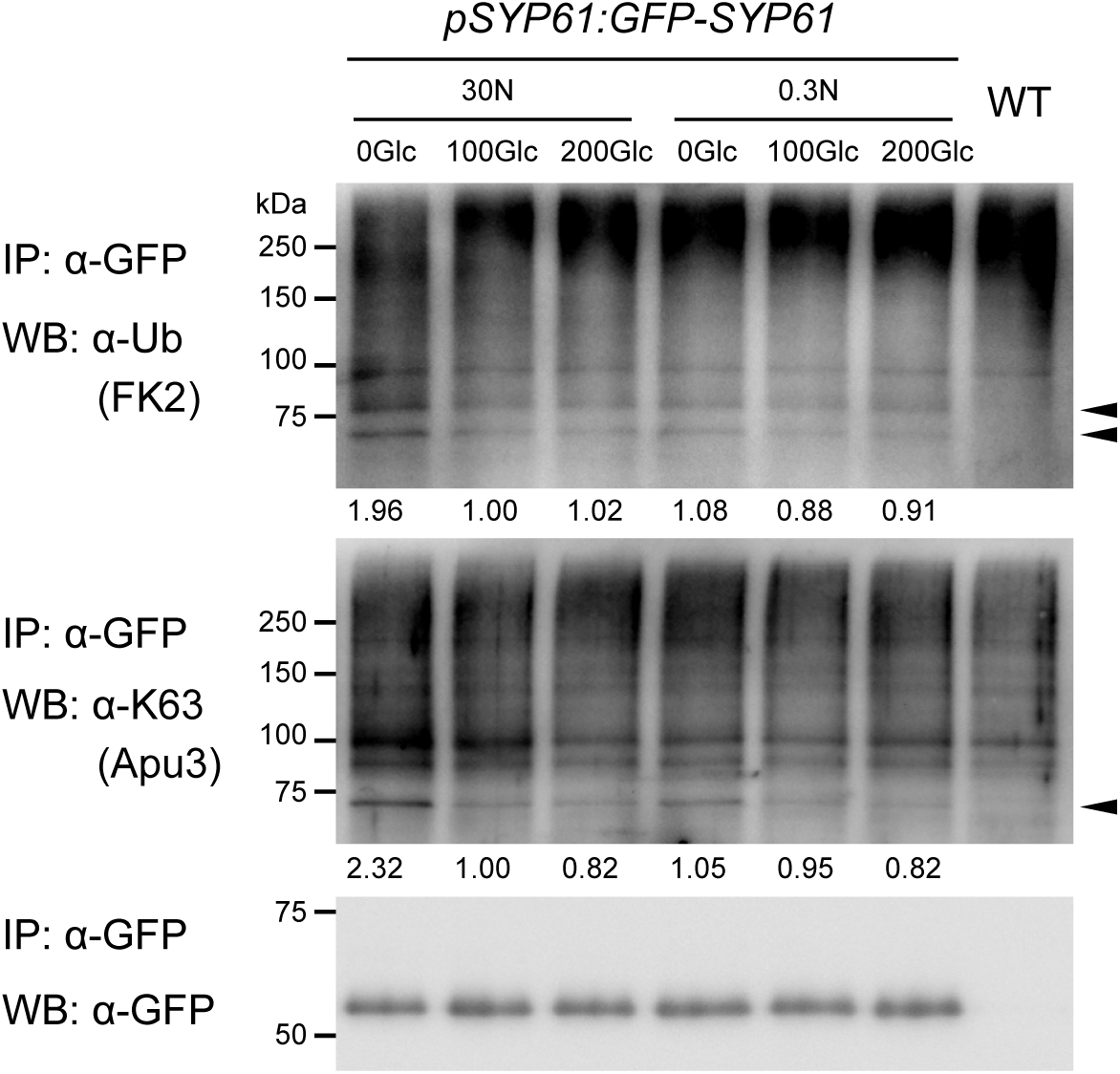
Ubiquitination of SYP61 is affected by C/N-nutrient availability. GFP-SYP61 (54.7 kDa) expressing Arabidopsis plants (*pSYP61:GFP-SYP61*) were grown for 10 days in 100 mM glucose/30 mM nitrogen containing liquid medium, and treated with the media containing indicated concentration of glucose and nitrogen for 3 h. Extracted proteins were immunoprecipitated with anti-GFP antibody beads, and detected with anti-GFP, anti-ubiquitin (FK2), and anti-K63-linked ubiquitin (Apu3) antibodies. Arrowheads indicate ubiquitinated GFP-SYP61. The numbers are showing the relative intensity of the di-Ub bands (the lower band indicated in anti-ubiquitin) normalized by the band intensity of GFP-SYP61. Glc, glucose; N, nitrogen; unit, mM. WT, wild type.

## Discussion

SNARE protein SYP61 is an important component of post-Golgi membrane trafficking(A. A. Sanderfoot et al., 2001; Uemura et al., 2004). SYP61 predominantly localized to the TGN/EE, but recent studies reported that SYP61 also localized to the plasma membrane and the vacuolar membranes under specific conditions(Gendre et al., 2013; Hachez et al., 2014; Rosquete et al., 2019). In addition, SYP61 colocalized with the LE markers, an R-SNARE VAMP727 and a RAB GTPase ARA7/RABF2b, to form the hybrid compartment of the TGN/EE and the LE involved in the endocytosis of FLS2 receptor protein after flg22 treatment(Choi et al., 2013). This promiscuous behavior of SYP61 suggests an as yet unknown post-translational mechanism involved in the regulation of SYP61 localization and function in response to environmental stimuli. In this study, we found that SYP61 is ubiquitinated in plants with K63-linked ubiquitin chain; to our knowledge, this is the first report of post-translational modification of SYP61. Moreover, the ubiquitination including the K63-linked chain was affected by the plant C/N-nutrient availability. Recent studies revealed several functions of ubiquitination on SNARE proteins in yeast and mammals. For example, mono-ubiquitination and deubiquitination of human Golgi-localized SNARE protein syntaxin5 (Syn5) regulate SNARE complex formation during mitosis and mediates Golgi structure reassembly(Huang, Tang, & Wang, 2016). K63-linked ubiquitination of the yeast SNARE protein Snc1 resulted in its direct binding to COPI, allowing it to be trafficked back from the endocytic to the exocytic pathway(Xu et al., 2017). In plants, a plasma membrane localized SNARE SYP122 was detected in the K63-linked ubiquitome analysis(Johnson & Vert, 2016), but its upstream and downstream is not revealed.

We also demonstrated that the ubiquitin ligase ATL31 catalyzes K63-linked ubiquitination of SYP61 *in vitro* and in plant cells. Mass spectrometry analysis identified several sites of SYP61, especially in the SNARE domain, directly ubiquitinated by ATL31. This modification would be an obstacle to interact with other SNARE proteins, therefore it may inhibit SYP61 to form a SNARE complex with inappropriate SNARE partners and timing. Our finding that the *syp61* knockdown mutants were hypersensitive to disrupted C/N-nutrient stresses, with a phenotype similar to that of the *atl31* loss-of function mutant(Sato et al., 2009), is suggesting the cooperative function of ATL31 and SYP61. The function of the SYP61 ubiquitination by ATL31 remains an open question.

Our MS-based and immunoblotting analyses indicated that ATL31 catalyzes multiple ubiquitin chain types, suggesting that ATL31 mediates multiple cellular signals via ubiquitination of distinct target proteins. Our previous study reported that ATL31 targets 14-3-3 proteins and subjects to proteasomal degradation under C/N-nutrient stress conditions(Sato et al., 2011; Yasuda et al., 2014). ATL31 is phosphorylated by CIPK7/12/14 kinases under C/N-nutrient stress, and ATL31 interacts with 14-3-3 through this phosphorylation(Yasuda et al., 2017). Thus, the activity of ATL31, including the target interactions and possibly the type of ubiquitin linkages, seems to be tightly regulated in response to environmental conditions. The SYP61 ubiquitination by ATL31 might be also activated by as-yet-unknown upstream signals.

In addition to C/N-nutrient stress, ATL31 positively regulates plant resistance to pathogen attacks and possibly to salt stress(Maekawa et al., 2014, 2012; Peng et al., 2014). In plant immunity, ATL31 enhances callose deposition(Maekawa et al., 2014, 2012). ATL31 targeted to fungal penetration sites in association with SYP121/PEN1, and accelerated the formation of rigid cell walls called papillae(Maekawa et al., 2014), suggesting that ATL31 possibly mediates vesicle trafficking of cell wall material and/or the related enzymes. SYP61 is also reported to be involved in salt stress tolerances and cell wall secretion. *Osm1*, the knockdown mutant of SYP61 is hypersensitive to the salt and osmotic stresses(Zhu et al., 2002). Proteomics on SYP61 compartment has identified cell wall relating enzymes as potential cargos(Drakakaki et al., 2012), and recent advanced glycomic analysis reported that this mutant also shows modulated trafficking of cell wall materials, such as polysaccharides and glycoproteins(Wilkop et al., 2019). These functional similarities of ATL31 and SYP61 suggest their possible relevance in diverse physiological processes in plants.

In this report, we demonstrated the involvement of a SNARE protein SYP61 and its ubiquitination in plant C/N-nutrient responses. Our findings provided new insights into the ubiquitin signaling and membrane trafficking machinery in plant.

## Methods

### Plant material and growth conditions

The *Arabidopsis thaliana* ecotypes Columbia-0 (Col-0) and C24 were used in this study. The T-DNA insertion mutant *osm1*(Zhu et al., 2002) was kindly provided by Dr. Jianhua Zhu (University of Maryland). Transgenic Arabidopsis plants, *p35S:ATL31-GFP, p35S:ATL31*^*C143S*^*-GFP, pSYP61:mRFP-SYP61, pSYP43:mRFP-SYP43, pVHAa1:VHAa1-mRFP, pARA7:mRFP-ARA7* and *pVAMP727:tagRFP-VAMP727* have been described(Sato et al., 2009; Uemura, Ueda, & Nakano, 2012). Co-expression lines were generated by crossing and F1 lines were used for microscopic observation. For seed amplification, Arabidopsis seeds were surface-sterilized and sowed on 1x Murashige and Skoog (MS) medium supplemented with 1% sucrose and vitamins (pH5.7). After kept in dark at 4°C for 2-4 days to synchronize germination, the plants were grown under 16 h light/8 h dark at 22°C on the plates for several weeks, and transferred to flowerpot with soil containing compost and vermiculate in the ratio of 1:6. Tap water was used for watering. The date of the starting growth is counted as 0 DAS (days after stratification) in this paper. *Nicotiana benthamiana* plants were used for transient protein expression. The surface-sterilized seeds were sowed on 1xMurashige and Skoog (MS) medium supplemented with 1% sucrose and vitamins (pH5.7), and grown under 16 h light/8 h dark at 22°C on the plates for two weeks, then transferred to flowerpot with soil containing compost and vermiculate in the ratio of 1:6. Tap water was used for watering, and supplied with 1/1000 diluted Hyponex (HYPONeX JAPAN CORP, Japan) once a week.

For microscopy imaging of Arabidopsis root cells, seeds were vertically grown on modified Murashige and Skoog (MS) medium containing 100 mM glucose, 30 mM nitrogen (10 mM NH_4_NO_3_ and 10 mM KNO_3_), vitamins and 0.8% agar (pH5.7). The preparation protocol of the modified MS medium has been described(Sato et al., 2009). For vacuolar accumulation analysis, *p35S:ATL31-GFP* seeds were grown as described for 6 days, and on the 7^th^ day, the seedlings were exposed to light for 3 h before being wrapped in a double layer of aluminum foil and incubated for 10 h in the dark.

For the detection of the ubiquitination of SYP61 in Arabidopsis, seeds were grown in liquid 1xMS medium supplemented with 1% sucrose and vitamins (pH5.7) for 10 days under constant light with shaking at 70 rpm.

To test the effect of C/N-nutrient condition on SYP61 ubiquitination, seeds were grown in modified MS liquid medium containing 100 mM glucose, 30 mM nitrogen (10 mM NH_4_NO_3_ and 10 mM KNO_3_), vitamins and 2 mM MES (pH5.7) for 10 days under constant light with shaking at 70 rpm, and treated with various MS media containing different concentration of glucose, nitrogen and mannitol for 3 h.

### Plant transformation

To generate transgenic plants expressing GFP-SYP61, translational fusions between cDNAs for GFP and SYP61 were generated using fluorescent tagging of full-length proteins method(Tian et al., 2004). Briefly, the cDNA encoding GFP was inserted in front of the start codon of the *SYP61* gene. These constructs included 2.0 kb of the 5’-flanking sequence and 0.8 kb of the 3’-flanking sequence of SYP61. The chimeric fragments were subcloned into the binary vector, pGWB1(Nakagawa et al., 2007), followed by transformation of Arabidopsis plants.

The amiRNA used was designed to target SYP61 using the online WMD interface (http://wmd2.weigelworld.org)(Ossowski, Schwab, & Weigel, 2008; Schwab, Ossowski, Riester, Warthmann, & Weigel, 2006). The amiRNA construct was amplified by PCR, using the primers listed in Supplementary Table S2 and the plasmid pRS300(Ossowski et al., 2008) as template. The PCR product was subcloned into the pGWB402-omega vector(Nakagawa et al., 2007). To generate transgenic Arabidopsis plants, these constructs were introduced into *Agrobacterium tumefaciens* GV3101 (pMP90) by electroporation, followed by transformation of Arabidopsis plants (Col-0) by the floral dipping method(Clough & Bent, 1998).

The primers used in these experiments are listed in Supplementary Table S3.

### Transient expression in *Nicotiana benthamiana*

For the plasmid construction, coding sequences of each gene were amplified by PCR and cloned into vector pENTR/D-TOPO (Life Technologies). These fragments were subsequently transferred to destination vectors using the Gateway system according to the manufacturer’s protocol (Invitrogen). For immunoprecipitation assays, coding sequences of *ATL31* and *ATL31*^*C143S*^ were subcloned into the pGWB11 destination vector, *UBQ1* (At3g52590) was subcloned into pEarleygate203(Earley et al., 2006), and SYP61 (At1g28490) was subcloned into pGWB6 and pGWB12 destination vectors(Nakagawa et al., 2007). For the *in vivo* ubiquitination analysis, coding sequences of ATL31 and ATL31^C143S^ were transferred into the pAMPAT-GW-3xFLAG destination vector(Yamada et al., 2016), and using them as templates, the sequences from ATL31 or ATL31^C143S^ to 3xFLAG were subcloned into the pGWB502-Ω destination vector(Nakagawa et al., 2007). All constructs were introduced into *Agrobacterium tumefaciens* strain GV3101 (pMP90) by electroporation. The *A. tumefaciens* containing the constructs were grown in 2xYT liquid medium at 28°C overnight with shaking, and the cells were resuspended with infiltration buffer (10 mM MES, 10 mM MgCl_2_, and 450 µM acetosyringone, pH 5.6). The mixtures of the suspension were infiltrated into the leaves of 5-6-week-old *N. benthamiana* by syringe without needle. The *A. tumefaciens* carrying p19 suppressor was co-infiltrated with the all samples(Takeda et al., 2002). After 3 days, the leaves were harvested with liquid nitrogen.

### Protoplast transfection

ATL31 was subcloned into the vector pUGW5(Nakagawa et al., 2007), ST-mRFP was subcloned into pBS, and mRFP-SYP61 and mRFP-VAMP727 were subcloned into the pUC vectors as described before(Uemura et al., 2004). The plasmids were introduced by polyethylene glycol-mediated transformation(Yoo, Cho, & Sheen, 2007) into Arabidopsis mesophyll protoplasts prepared from leaf tissues, and images were taken 16 h later.

### Immunoprecipitation

The plant materials were frozen with liquid nitrogen, and powdered using morter and pestle, and the expressed proteins were extracted using protein extraction buffer (50 mM Tris, 0.5% Triton X-100, 150 mM NaCl, 10% glycerol, 1 mM EDTA, pH 7.5) supplemented with 10 µM MG132 and Complete Protease Inhibitor Mixture (Roche Applied Science). To detect ubiquitination, the deubiquitination inhibitors 20 mM N-ethylmaleimide and 2 mM 1,10-phenanthroline were added. To test the *in vivo* ubiquitination of SYP61 by ATL31, and the effect of C/N-nutrient conditions, 500 nM of MLN7243 (Active Biochem, A-1384) was also added as an E1 inhibitor to avoid additional ubiquitination during the extraction steps. The mixtures were centrifuged at 20,000x*g* for 5 min at 4°C and the supernatants were collected. This step was repeated twice, and then the proteins were immunoprecipitated with anti-FLAG M2 affinity gel (Sigma-Aldrich, M8823), anti-GFP mAb-magnetic agarose (MBL, D153-10) or Anti-GFP mAb-Magnetic Beads (MBL, D153-11). D153-11 was used for checking ubiquitination of GFP-SYP61 in response to C/N-nutrient conditions, and D153-10 was used for other experiments to immunoprecipitate GFP-tagged proteins. The extracts and the beads were mixed and rotated at 4°C for 1 h, and washed for three or four times with the extraction buffer without inhibitors. To detect FLAG-SYP61 ubiquitination, proteins were eluted with 150 μg.ml^-1^ 3× FLAG peptide (Sigma-Aldrich, F4799) in extraction buffer, followed by precipitation in cold acetone, resuspension in SDS sample buffer (62.5 mM Tris, 2% SDS, 10% glycerol, 5% 2-mercaptoethanol, 0.01% bromophenol blue, pH 6.8) and incubation at 90°C for 5 min. In other experiments, proteins were eluted with the SDS sample buffer at 90°C for 5 min or 55°C for 30 min.

Proteins were separated by SDS-PAGE and detected by immunoblotting with anti-FLAG (Wako, 018-22381), anti-SYP61, anti-GFP (MBL, 598), anti-ubiquitin (FK2) (Wako, 302-06751), anti-K48 ubiquitination (Apu2) (Millipore, 05-1307) and anti-K63 ubiquitination (Apu3) (Millipore, 05-1308) antibodies. Anti-SYP61 antibody was generated in a rabbit by immunization with recombinant GST-SYP61. GST-tagged truncated SYP61, which included amino acid residues 1 to 218, was expressed in *E. coli* Rosetta-gami (DE3), purified and injected into a rabbit to generate an anti-SYP61 polyclonal antibody. The antibody was purified by protein G affinity column chromatography (GE Healthcare) (Supplementary Fig. S5).

### Split ubiquitin yeast two-hybrid assay

All assays used the yeast strain L40ccua (*MATa his3Δ200 trp1-901 leu2-3,112 LYS2::*(lexAop)_4_*-HIS3 ura3::*(lexAop)_8_*-lacZ ADE2::*(lexAop)_8_*-URA3 gal80* can^R^ cyh2^R^). The full-length coding sequences of *ATL31* and *ATL31*^*C143S*^ were subcloned into the destination vector pMetYC_GW, containing the C-terminal half of ubiquitin, and the full-length coding sequence of *SYP61* was subcloned into the destination vector pNX32_GW, containing the N-terminal half of ubiquitin(Obrdlik et al., 2004). The empty vector pNX32_GW was used as a negative control. Constructs were transfected into yeast using Frozen-EZ Yeast Transformation II Kits (Zymo Research) according to the manufacturer’s protocol, with growth assessed as described in the Yeast Protocols Handbook (Clontech).

### *In vitro* ubiquitination assay

*In vitro* ubiquitination assays were performed as described(Sato et al., 2009). Briefly, 500 ng of GST-SYP61 or GST were incubated with 500 ng of MBP-ATL31 for 0-3 hours at 30°C in 30 µL of reaction mixture, containing 50 ng E1 (Wako, 219-01111), 250 ng UbcH5a for E2 (Wako, 215-01191), 9 µg ubiquitin (Sigma-Aldrich, U6253), 25 µM MG132, 40 mM Tris-HCl (pH 7.5), 5 mM MgCl_2_, 2 mM ATP and 2 mM DTT. Reactions were stopped by adding SDS sample buffer. Proteins were separated by SDS-PAGE and detected by immunoblotting using anti-MBP (New England BioLabs, E8032S), anti-GST (MBL, M071-3), anti-ubiquitin (FK2) (Wako, 302-06751), anti-K48 ubiquitination (Apu2) (Millipore, 05-1307) and anti-K63 ubiquitination (Apu3) (Millipore, 05-1308) antibodies.

### Mass spectrometry analysis of ubiquitination sites

Ubiquitinated GST-SYP61 proteins were separated by SDS-PAGE and stained with SYPRO Ruby (Lonza, Switzerland) as described in the manufacturer’s protocol. Gel pieces containing ubiquitinated GST-SYP61 were excised, dehydrated with 100% acetonitrile and incubated in 10 mM dithiothreitol and 50 mM ammonium bicarbonate for 45 min at 56°C with shaking. The gel pieces were subsequently incubated in 55 mM chloroacetamide/50 mM ammonium bicarbonate for 30 min at room temperature, washed with 25 mM ammonium bicarbonate and dehydrated with 100% acetonitrile. The dried gels were incubated for 16 h at 37°C in 50 mM ammonium bicarbonate containing sequence grade modified trypsin (Trypsin Gold; Promega, USA). The digested peptides were eluted from the gels with 50% acetonitrile (v/v) /5% formic acid (v/v) and dried using an evaporator. The peptides were dissolved in 2% acetonitrile (v/v) /0.1% formic acid (v/v) and filtered with Ultrafree-MC Centrifugal Filters (PVDF 0.45 μm; Millipore, USA). Peptides were identified using an EASY-nLC 1000 liquid chromatograph coupled to an Orbitrap Mass Spectrometer (Thermo Scientific, USA), followed by assessment using a SEQUEST algorithm embedded in Proteome Discoverer 1.4 software (Thermo Scientific, USA) against TAIR10 (http://www.arabidopsis.org/index.jsp), as described(Lu et al., 2016).

### Gene expression analysis

Total RNA was isolated from the 7-days-grown Arabidopsis seedlings using TRIzol reagent (Invitrogen), and was treated with RQ1 RNase-free DNase (Promega) according to the manufacturer’s protocols. Then, the cDNA was synthesized using oligo(dT) primer (Promega) with 18S rRNA specific primer and ReverTraAce reverse transcriptase (Toyobo). qRT–PCR analysis was performed using SYBR premix Ex Taq (TaKaRa) and Mx3000P (Agilent Technologies) according to the manufacture’s protocol. The used primers are listed in Supplementary Table S4.

### Carbon/nitrogen-nutrient response analysis

Seeds of each genotype were surface-sterilized and sown on modified MS-based solid medium containing glucose, mannitol and nitrogen (KNO_3_ and NH_4_NO_3_ were used in 1:1 molar ratio) at the concentration indicated in figures. The medium was prepared as described(Sato et al., 2009). After kept in dark at 4°C for 2 days to synchronize germination, the plants were grown under 16 h light/8 h dark at 22°C. The percentage of the seeds with green spread cotyledons in the germinated seeds were calculated.

### Chemical treatment

5 or 6 day-old seedlings were incubated in 1 mL of modified liquid MS medium 100 mM glucose/30 mM nitrogen containing 50 μM Brefeldin A (BFA), or 33 μM Wortmannin (WM). Fluorescence was monitored by confocal microscopy 30 min and 1 h later respectively. Seedlings treated with BFA were pre-incubated in 5 μM FM4-64 for 15 min and then transferred to BFA containing medium.

### Confocal laser-scanning microscopy

Confocal laser-scanning microscopy was performed using Zeiss LSM510 and LSM780 microscopes. GFP fluorescence was excited by a 488 nm argon laser and detected using a 505–550 nm band-pass emission filter, whereas RFP fluorescence was excited by a 561 nm DPSS laser and detected using a 575–615 nm band-pass emission filter. FM4-64 fluorescence was excited by a 561 nm DPSS laser and detected using a 575 nm long-pass emission filter. Images were processed using ImageJ software (National Institutes of Health, Maryland, Washington, DC).

### Quantitative ubiquitin chain type analysis with AQUA peptide

MS/MS-based absolute quantitation (AQUA) of ubiquitin peptides by parallel reaction monitoring (PRM) was performed as described previously(Tsuchiya et al., 2017). Proteins were separated by a short run (1 cm) on 4–12% NuPAGE gels, and the gel region corresponding to molecular weight >75 kDa was subjected to trypsinization. Trypsinized peptides were extracted, spiked with ubiquitin AQUA peptides, and analyzed in targeted MS/MS mode on a Q Exactive mass spectrometer coupled to an EASY-nLC 1000 (Thermo Scientific). Raw data were processed using the PinPoint software version 1.3 (Thermo Scientific).

### Statistics

All values shown as bar graphs are mean ± s.d or s.e.m as stated. *P* values were calculated by two-tailed Welch’s *t*-test for two-group comparisons, and by two-tailed Dunnett’s test for multiple-group comparisons. Statistical significance was set based on *P*-values. n.s., *P* > 0.05; \**P* < 0.05; \*\**P* < 0.01

## Acknowledgments

We thank Drs. Tsuyoshi Nakagawa (Shimane University, Japan) and Yoshihisa Ueno (Ryukoku University, Japan) for kindly providing the Gateway destination vectors used in this study. We also thank Drs. Bernhard Grimm (Humboldt University, Germany) and Ryouich Tanaka (Hokkaido University, Japan) for kindly providing materials for split ubiquitin yeast two-hybrid assays. The *osm1* mutant seeds were kindly provided by Dr. Jianhua Zhu (University of Maryland, USA). We thank Dr. Ooi-Kock Teh (Hokkaido University, Japan) for kind support in microscope analysis, and the Instrumental Analysis Division, Global Facility Center, Creative Research Institution, Hokkaido University for mass spectrometry analysis. This work was supported by a Grant in-Aid for Scientific Research to T.S. [No. 17K08190 and 20K05949] from the Japan Society for the Promotion of Science (JSPS), by a grant from the NOASTEC foundation, Hokkaido University Young Scientist Support Program to T.S., and by Grants in Aid for Scientific Research to J.Y. [Nos. 15H0116705, 2629218885, and 2666004604] from the JSPS. R.P.B. was funded by grant from Knut and Alice Wallenberg Foundation. Y.H. is a recipient of JSPS Research Fellowship for Young Scientists. T.H.R. was supported by MEXT Honors Scholarship for Privately Financed International Students.

## Author contributions

Y.H. and T.S. designed the experiments. Y.H., T.H.R., T.U., A.F., Y.L., Y.M., Y.S., A.B., S.M., S.Y., K.M, and Y. F. performed the experiments. Y.H., Y.S., K.T., A.N., R.P.B. and T.S. analyzed the data. Y.H., T.H.R. J.Y. and T.S. wrote the article, and Y.S. and R.P.B. edited it. All the authors approved and edited the manuscript.

