## Supplementary Figures and Tables for "TGN/EE SNARE protein SYP61 is ubiquitinated and required for carbon/nitrogen-nutrient responses in Arabidopsis"

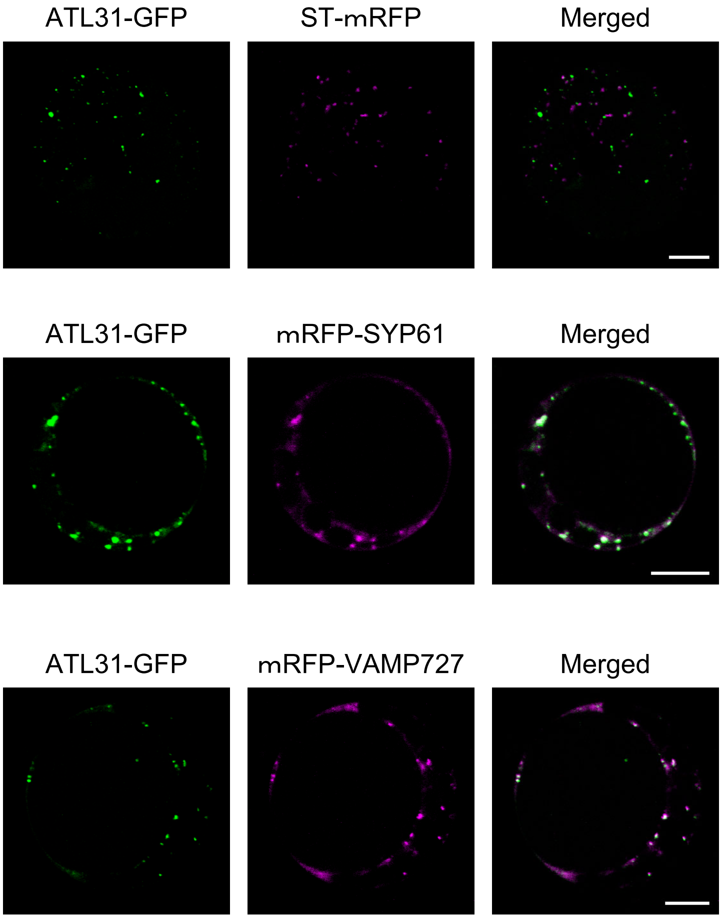


**Supplementary Fig. S1 Co-localization analysis of ATL31 with endosome markers.** Representative confocal images of Arabidopsis mesophyll protoplast cells transiently co-expressing ATL31-GFP with ST-mRFP (Golgi), mRFP-SYP61 (TGN/EE), and mRFP-VAMP727 (late endosome). Bars = 10 μm.

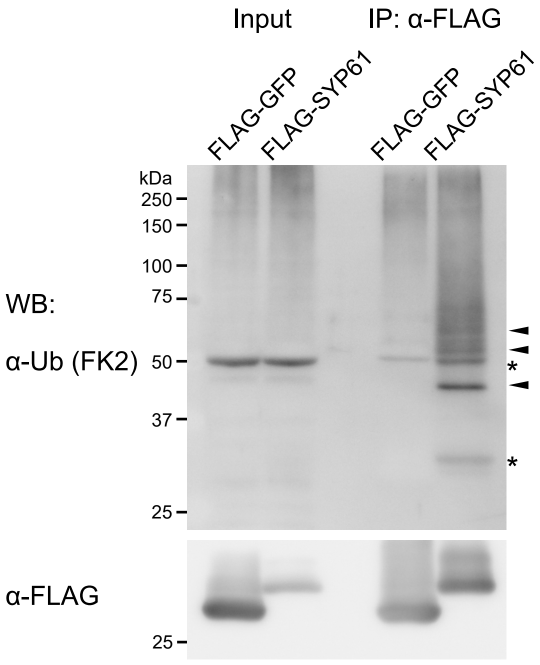


**Supplementary Fig. S2 SYP61 is ubiquitinated in plant.**

FLAG-GFP (30 kDa) or FLAG-SYP61 (30.7 kDa) was transiently expressed in *N. benthamiana* leaves. Extracted proteins were immunoprecipitated with anti-FLAG antibody beads, and detected with anti-FLAG and anti-ubiquitin (FK2) antibodies. Arrowheads indicate ubiquitinated SYP61, and asterisks indicate unknown bands.

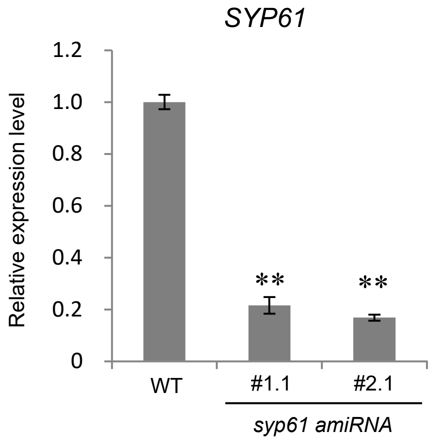


**Supplementary Fig. S3 Generating *syp61 amiRNA* knockdown plants.**

Relative expression levels of of *SYP61* mRNA transcript normalized by *18S rRNA*. Total RNA was extracted from WT (Col-0) and *syp61* knockdown mutant (*syp61 amiRNA*) seedlings grown for 7 days. Data are means ± s.e.m (n = 4). Asterisks indicate significant differences compared with the WT as determined by Dunnett′s test (***P*<0.01).

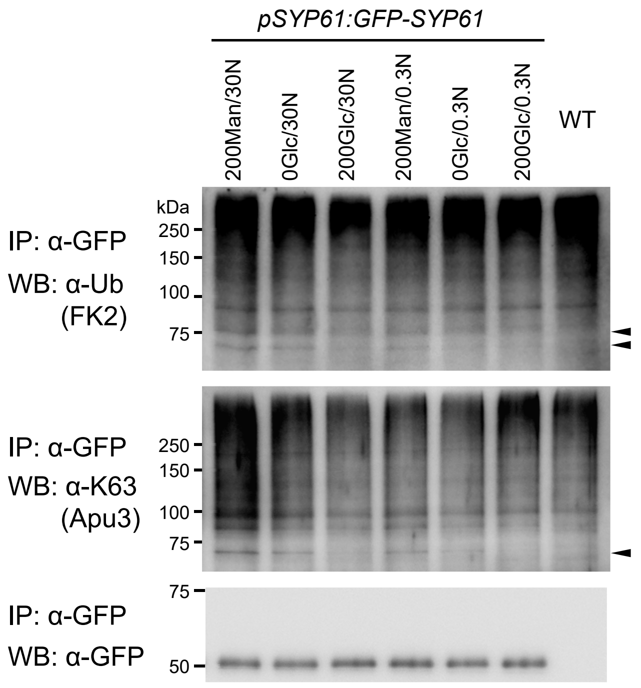


**Supplementary Fig. S4 Ubiquitination of SYP61 is affected by C/N-nutrient availability.**

GFP-SYP61 (54.7 kDa) expressing Arabidopsis plants (*pSYP61:GFP-SYP61*) were grown for 10 days in 100 mM glucose/30 mM nitrogen containing liquid medium, and treated with the media containing indicated concentration of mannitol, glucose and nitrogen for 3 h. Extracted proteins were immunoprecipitated with anti-GFP antibody beads, and detected with anti-GFP, anti-ubiquitin (FK2), and anti-K63-linked ubiquitin (Apu3) antibodies. Arrowheads indicate ubiquitinated GFP-SYP61. Man, mannitol; Glc, glucose; N, nitrogen; unit, mM.

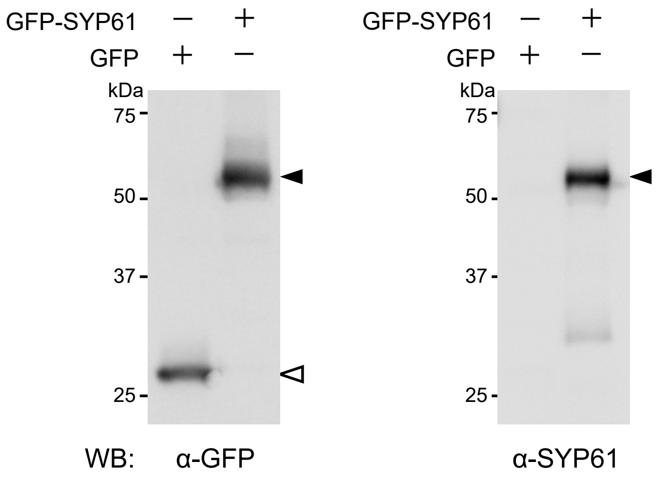


**Supplementary Fig. S5 Recognition of SYP61 protein by anti-SYP61 antibody.**

Proteins were extracted from *N. benthamiana* leaves transiently expressing GFP (27 kDa) or GFP-SYP61 (56.2 kDa), and detected with anti-GFP and anti-SYP61 antibodies. The same amount of proteins were applied. Closed arrowheads indicate GFP-SYP61, and open arrowhead indicates GFP.

**Supplementary Table S1.**
**Ubiquitinated SYP61 peptides detected by mass spectrometry analysis.**

| Peptide sequence[a] | Position | q-Value[b] | XCorr[b] |
|---|---|---|---|
| SGVLAG**K**VSSGAGHASEVR | K106 | 0 | 3.21 |
| QMLLI**K**QQDEELDELSK | K157 | 0 | 1.94 |
| QQDEELDELS**K**SVQR | K168 | 0 | 2.39 |
| IIDELDTEMDST**K**NR | K202 | 0 | 3.50 |
| IIDELDTEMDST**K**NRLEFVQK | K202 | 0 | 2.45 |
| NRLEFVQ**K**K | K210 | 0 | 2.76 |

[a]Ubiquitinated lysine (K) residue is indicated by bold letter with underline.
[b]The scores assigned by SEQUEST software after database searching.

**Supplementary Table S2. Primer sequences used for *syp61 amiRNA* construction.**

| Primer name | Sequence (5'→3') |
|---|---|
| Primer I* | gaTCATTAGTTCTCTACGCGCTTtctctcttttgtattcc |
| Primer II* | gaAAGCGCGTAGAGAACTAATGAtcaaagagaatcaatga |
| Primer III* | gaAAACGCGTAGAGATCTAATGTtcacaggtcgtgatat |
| Primer IV* | gaACATTAGATCTCTACGCGTTTtctacatatatattcct |
| pRS300_A | CTGCAAGGCGATTAAGTTGGG |
| pRS300_B | CTGTTTCCTGTGTGAAATTGTTATCCGC |

*The regions encoding microRNA specific to *SYP61* mRNA are shown in capital letters.

**Supplementary Table S3. Primer sequences used for the plasmid construction.**

| Gene | Sequence (Forward, 5'→3') | Sequence (Reverse, 5'→3') |
|---|---|---|
| *SYP61* | CACCATGTCTTCAGCTCAAGA | GGTCAAGAAGACAAGAACGAA |
| *UBQ1* | CACCATGCAGATCTTCGTGAAA | CTAACCTCCTCTAAGCCTCAACAC |
| *ATL31-3xFLAG* | CACCATGGATCCCATAAAACACAT | CTAAGATCTCTTGTCATCGTCATC |

**Supplementary Table S4. Primer sequences used for expression analysis.**

| Gene | Sequence (Forward, 5'→3') | Sequence (Reverse, 5'→3') |
|---|---|---|
| *SYP61* | GATTCAGGATTCTATTGATAAGTTGC | CTCATCTACCTGCCACTCAATG |
| *18S rRNA* | CGGCTACCACATCCAAGGAA | GCTGGAATTACCGCGGCT |
